# Pressure drives rapid burst-like collective migration from 3D cancer aggregates

**DOI:** 10.1101/2021.04.25.441311

**Authors:** Swetha Raghuraman, Ann-Sophie Schubert, Stephan Bröker, Alejandro Jurado, Annika Müller, Matthias Brandt, Bart E. Vos, Arne D. Hofemeier, Fatemeh Abbasi, Martin Stehling, Raphael Wittkowski, Timo Betz

## Abstract

Collective migration of cells is a key behaviour observed during morphogenesis, wound healing and cancer cell invasion. Hence, understanding the different aspects of collective migration is at the core of further progress in describing and treating cancer and other pathological defects. The standard dogma in cell migration is that cells exert forces on the environment to move and cell-cell adhesion-based forces provide the coordination for collective migration. Here, we report a new collective migration mechanism that is independent of pulling forces on the extra-cellular matrix (ECM), as it is driven by the pressure difference generated inside model tumours. We observe a striking collective migration phenotype, where a rapid burst-like stream of HeLa cervical cancer cells emerges from the 3D aggregate embedded in matrices with low collagen concentration (0.5 mg ml^*−*1^). This invasion-like behaviour is recorded within 8 hours post embedding (hpe), and is characterised by high cell velocity and super-diffusive collective motion. We show that cellular swelling, triggered by the soft matrix, leads to a rise in intrinsic pressure, which eventually drives an invasion-like phenotype of HeLa cancer aggregates. These dynamic observations provide new evidence that pressure-driven effects need to be considered for a complete description of the mechanical forces involved in collective migration and invasion.

---

The mechanical interplay between cells and the ECM contributes vastly to the phenotypes of cell migration^1, 2^. While migrating independently or within solid tissues, cells constantly experience shear forces, compression, tension and hydrostatic as well as osmotic pressures^3–7^. Mechanical homeostasis ensures a complete force balance, such that only marginal motion of single cells within tissue is observed. However, this balance is broken in migrating cells that need to generate well-orchestrated forces, not only on the single-cell level but also on the collective level. Over the past decades, our understanding of cell migration has been boosted by a series of ground-breaking experiments that allowed not only for classifying different kinds of collective and single-cell migration, both *in vitro* and *in vivo*, but also for identifying a series of key molecular players driving this motion^1, 4, 8, 9^. Yet, our knowledge of forces and force generation is lagging vastly behind, with most insights suggesting that acto-myosin-originated forces solely lead to cell migration. This notion stands in contrast to a series of hallmark experiments in the field of tumour biophysics demonstrating that pressure, and pressure distribution within solid tumours can be critical for proliferation, migration and protein expression^10–16^. Summarising, these findings show that pressure is not only a consequence of cell proliferation in confined geometries, but may also play a pivotal role in cellular functions and tumour progression. Although it may seem intuitive that pressure relaxation could lead to collective cell migration, this has not been addressed so far as a realistic mechanism.

One reason this has not yet been investigated might be that in experimentally attractive *in vitro* systems of cancer spheroids embedded in ECM matrices, cells actively pull on the ECM, which can result in a release of pressure within the aggregate. In fact, cells at the periphery of *in vitro* spheroids, sense and respond to ECM via integrins, inducing changes in their morphology, polarity, and contractile forces by which they pull on the ECM^17–19^. For example, collective cell-generated forces have produced matrix contractile pressures of up to *≈*700 Pa, showing that, collectively, cells can strongly affect their environment^20^.

Here, we demonstrate that pressure-induced invasion is a collective migration mechanism that is fundamentally different from previously described adhesion-based strategies. Using a simple model system of HeLa cervical cancer aggregates embedded in 0.5 mg ml^*−*1^ collagen, cell swelling leads to a pressure increase that climaxes in a burst-like invasion as result of a series of mechanical events. After an initial contractile phase, where the cells potentially probe the mechanical environment of the soft ECM, cells undergo a volume increase that, consequently, raises the internal pressure. Then, rapid collective outbursts occur as the high pressures relax into regions with the least mechanical resistance. Using a computer simulation based on the initial experimental conditions, we can fully recapitulate the experimental result, further supporting the finding that pressure-driven collective invasion is a novel mechanical phenotype of cell motility.

## Rapid collective outbursts triggered by low density collagen microenvironment

To investigate pressure-driven collective migration we use cancer cell aggregates as standard model systems to study the interaction between 3D simple tissues and well controlled ECM microenvironments^17^ (Figure 1a). Exploiting light sheet-based 3D microscopy we followed the motion of individual HeLa cells within aggregates by tracking the fluorescently marked nuclei via histone H2B-mCherry or H2B-RFP. HeLa tumour models embedded in 2.5 mg ml^*−*1^ collagen I (higher concentration collagen, HCC) did not show any shape changes but simply grew due to proliferation (Figure 1b). However, by merely reducing the collagen concentration to 0.5 mg ml^*−*1^ (lower collagen concentration, LCC) without any other changes to the cell culture or sample preparation, we observed a drastic migration phenotype marked by rapid cell outbursts starting between 6-8 hpe, in which a large amount of cells were expelled into the surrounding ECM (Figure 1b, e, Supplementary Videos 1, 2).

**Figure 1:**
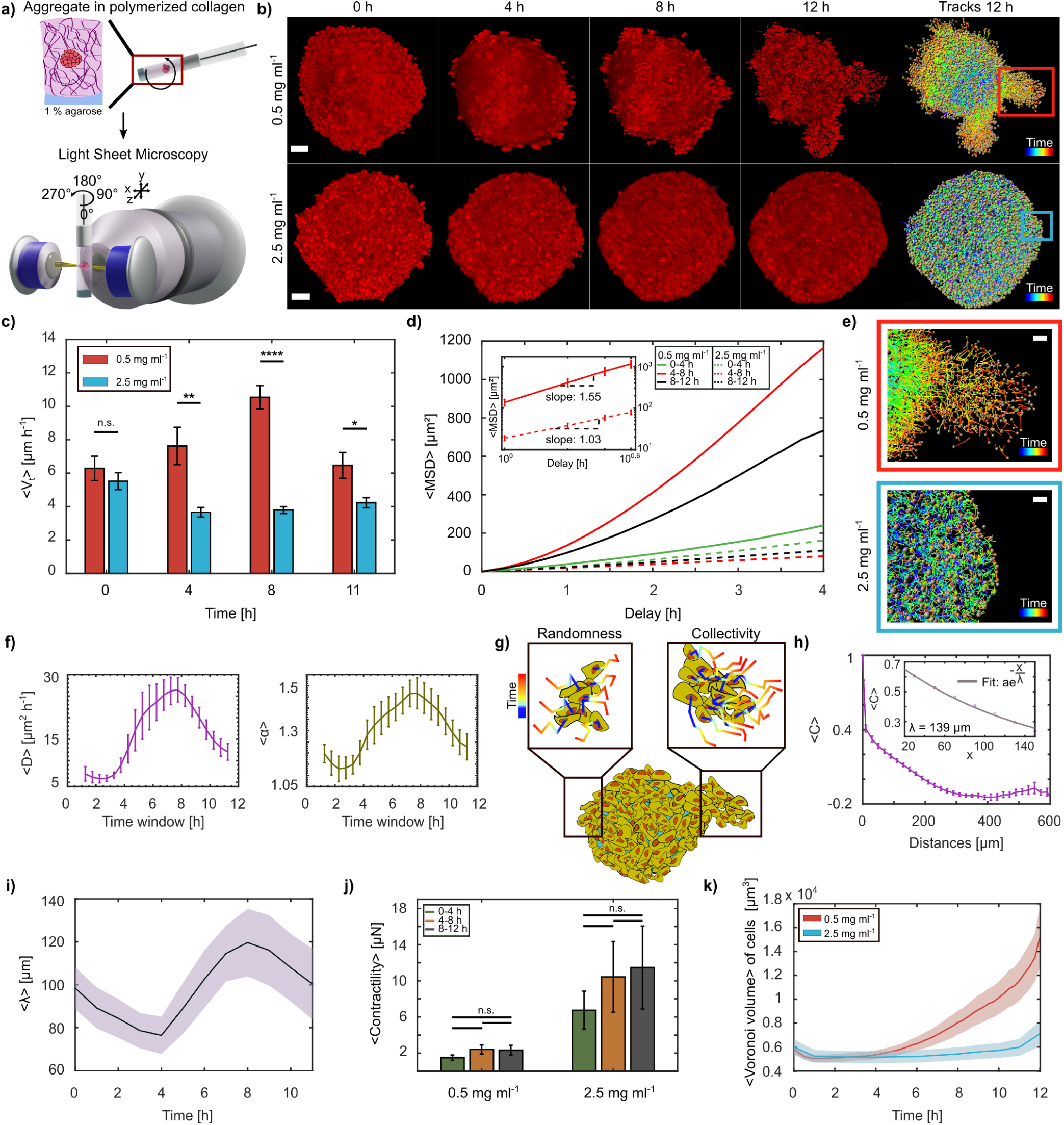
HeLa aggregates collectively burst in 0.5 mg ml^*−*1^ collagen matrix. **a)** Schematic representation of aggregate preparation and imaging process. **b)** Over 12 h, aggregates in 0.5 mg ml^*−*1^ collagen (LCC; top) burst into the matrix, whereas aggregates embedded in 2.5 mg ml^*−*1^ collagen (HCC; bottom) did not. Nuclei tracks at 12 h from each condition are shown, scale bars = 50 *µ*m. **c)** Comparison of average absolute cell velocities ⟨*V*_t_⟩ within aggregates embedded in LCC & HCC at different time points (N = 10 each, standard error of the mean (s.e.m.)). **d)** Average MSD for three time windows (N = 10 each). Inset shows the MSD in log scale between 4-8 h for LCC and HCC, with their corresponding slopes (s.e.m.). **e)** Zoomed-in view of the bursts (red box) and non-invasive (blue box) regions from (b) (scale bars = 20 *µ*m). **f)** Averages of diffusivity (D) and powerlaw exponent *α* obtained by fitting *f(x)* = *α x* + 6*D* onto the log of window MSDs (N = 10 each, s.e.m.). **g)** Representation of both the early random and later collective migration as observed in LCC. **h)** Velocity correlation ⟨C⟩ binned over distances (N = 10, bin size = 20 *µ*m). Exponential decay function fit on distances up to 150 *µ*m at 8 h (inset). **i)** Mean velocity correlation length acquired over time (median smoothed, N = 10, s.e.m.). **j)** Contractile stresses exerted by aggregates in LCC and HCC on their ECM (N =5 each, s.e.m.). **k)** Mean Voronoi volume of cells within aggregate acquired over time (N = 10, s.e.m.).

Quantitative analysis of the cell tracks revealed that, while during the first hour after embedding the overall velocity of the cell motion within the spheroid was similar in the different collagen concentrations, we find a drastic increase of mean absolute cell velocity within aggregates embedded in the LCC matrices when compared to the HCC (Figure 1c). Additionally, the visually obtained phenotype suggested a more directional motion during the bursts. Indeed, detailed analysis of the mean squared displacement (MSD) demonstrates that both conditions followed a power law ⟨*x*(*τ*)^2^⟩ = 6 D *τ*^*α*^, where *D* is the diffusion constant and *τ* the lag-time. However, while cells in the HCC behave diffusive, as quantified by an exponent of *α*_HCC_ = 1.03, in the burst phase we saw super-diffusive motion where *α*_LCC_ = 1.55, suggesting a ballistic behaviour (Figure 1d, g). To understand if the LCC condition had an immediate or rather delayed effect on this directional movement, we created a time-resolved MSD by a 2.5 h sliding window (Figure 1f, Supplementary Figure 1). Both the time evolution of the diffusion coefficient and the power law exponent showed time dependence, with a peak at 7-8 hpe, which is consistent with the observed morphological change. Surprisingly, the power law exponent showed a complex switching behaviour. After an initial slight super-diffusive behaviour (*α*_LCC,0_ = 1.2) it first tended briefly towards normal diffusion, before drastically increasing to its peak value and finally decreasing again, ending close to the initial value. This suggests that the coordination of the cell movement changes several times within a rather short time span of only 12 h. To further confirm this, we checked the correlation length of the velocity correlation function (Figure 1g, h), which quantifies collective migration. Indeed, the correlation length *λ* showed a similar behaviour as the power law exponent, with an initial decrease that was followed by a monotonic increase up to a peak value of about 120 *µ*m at 8 hpe, before it again decayed (Figure 1i). These rapid and fundamental changes in migration behaviour are intriguing, as they are typically associated with differential gene expression, for example, in the context of epithelial-mesenchymal transition^21^.

As the observed switching back and forth between random and collective migration seems at odds with biology, we considered a rather physical explanation. Therefore, we checked the forces exerted by the cells on the collagen matrix, however, they were not significantly changing over time, but confirmed the expected increase for HCC (Figure 1j). This finding is puzzling, as it shows that rapid and collective cell migration is not accompanied by a significant increase in traction forces applied on the ECM. To understand what factors are driving the outbursts, we checked the volume of the cells in the spheroid based on a Voronoi tessellation that uses the nucleus as a proxy for the cell position. As presented in Figure 1k, we saw an impressive and rapid increase in cell volume within aggregates in the LCC condition, whereas in HCC, the volume remained unchanged over the same time span.

## Changes in migration modes are accompanied by aggregate shape changes and a volume increase

To further understand both the changes in the motility characteristics and the volume change we developed a second approach to quantify the aggregate shape and volume by using a mathematical decomposition into spherical harmonics (SHs). SHs allow to develop any radial surface function into a series of shapes with increasing complexity, where the relative contribution of each shape is quantified by a single parameter (Figure 2a, b). As expected, the aggregate volume obtained from the SHs closely followed the time evolution that was observed for the Voronoi tessellation (Figure 2c). However, a close inspection of the different degrees and their relative contributions gave an additional surprise. While in the first two hours, the actual shape of the aggregate resembled an oblate, as quantified by the second mode of the SHs, we saw a rapid decrease of this mode within 5 hours, followed by a drastic increase of the higher modes marking the burst phase (Figure 2a, b) starting at 7 hpe. Hence, up until 4 hpe the aggregates changed their initial oblate shape towards a sphere without any volume change. Then, a massive volume increase sets in, which we hypothesize effectively pushed cells outwards in the observed bursts. This is consistent with the observed super-diffusive motion and higher collectivity, as the cells moved in a coordinated way. To test whether individual cells embedded in the LCC condition would also show the observed volume increase, we dispersed isolated HeLa cells in the LCC condition and measured the change in volume. Consistent with the aggregate, we found a drastic increase in volume immediately after seeding (Figure 2d). Interestingly, the volume increase in the aggregate started later, which suggests that in the tumour model embedded in LCC condition, the contact with other cells partially stabilises the cell volume.

**Figure 2:**
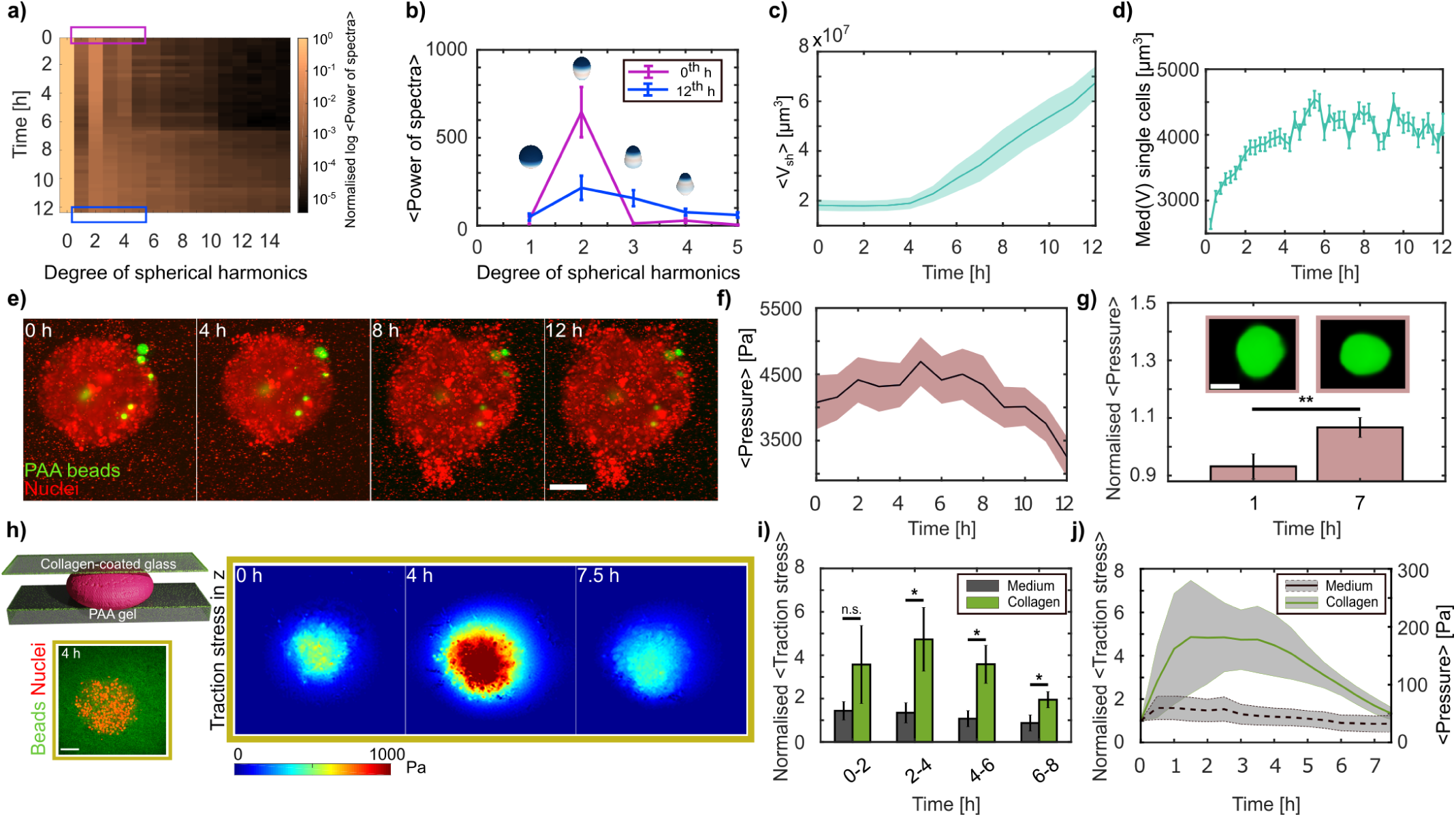
Cell volume increase leads to increased internal pressure. **a)** Temporal contribution of degrees 0-15 of spherical harmonics expansions on aggregate surfaces, colour coded by the normalised (by first degree) log of average power spectra (N = 10). **b)** Mean spherical harmonics spectra for hours 0 and 12, over the expansion degrees 1-5 (N = 10, s.e.m.). Scheme represents zero order shape contributions. **c)** Mean volume (⟨*V*_sh_⟩) increase as determined from the spherical harmonics (median smoothed, N = 10, s.e.m.). **d)** Median (Med) HeLa single cells volume *V* in LCC (n = 1694-2124 for each time point, N = 3, s.e.m.). **e)** Data showing PAA elastic beads (green) randomly distributed within aggregates (scale bar = 100 *µ*m). **f)** Mean anisotropic pressure recorded by polyacrylamide (PAA) beads (median smoothed). **g)** Anisotropic pressures normalised by the average pressure at the binned hours represented (bin size = 1 h). Insets depict deformation of one bead at 0 and 7 h (scale bar = 10 *µ*m). Statistical test was done on log normalised data. **h)** Schematic of sandwich experiment to confine aggregates and perform traction force microscopy (left, bottom: zoomed-in view of single plane image at the interface between PAA gel and the aggregate). Indicative data showing average traction stress in Pascals in z direction over time (right). Pushing (red) forces indicated. **i)** Normalised (by time 0) average traction stresses in z direction of aggregates confined with collagen embedding as compared to only medium. Statistical tests were done on log normalised data. **j)** Normalised (by time 0) and relative average aggregate pressure by the sandwich method. **(f, g)** n = 35-51, s.e.m.; **(i, j)** N = 6 (collagen), 5 (medium), s.e.m.

## Rise in internal aggregate pressure before burst phase

The observed lack of correlation between the ECM pulling forces and the outbursts suggests that the cells did not migrate out in a classical collective migration mechanism, but that the observed bursts were a collective effect due to the cell volume increase. Such a swelling may lead to a pressure increase inside the aggregate, similar to previous observations in mouse embryos and epithelial monolayers^5, 6^. To confirm such a volume-increase based pressure-rise hypothesis, we exploited recently introduced elastic hydrogel beads as force and pressure sensors, and analysed their deformation over the time course of the experiment (Figure 2e, Supplementary Video 3). Indeed, we found that the internal forces acting on the pressure sensors increased monotonically and showed a significant peak at 6-7 hpe, correlating with the onset of the outbursts (Figure 2f, g). After verifying that the pressure inside the aggregate was increasing, we wondered whether this could also exert a consequential pushing force on the environment, which would directly prove that the cells may be pushed out. To test this, we devised a modified 2.5D traction force microscopy approach^22, 23^, where the aggregate was confined in a sandwich-like fashion between a bottom polyacrylamide (PAA) gel of *Young’s modulus E ≈*1800 Pa and a top layer of collagen I-coated glass (Figure 2h), without compressing the aggregate. When the aggregates were surrounded by LCC, we could see a significant increase in pushing forces, marked by z-traction stress at 2-6 hpe of up to *≈*200 Pa, thereby deforming the bottom gel (Figure 2i, j). As the stiffness of the LCC was only about 10 Pa (confirmed by rheometry and from previous work)^24^, we did not expect that the aggregate would generate larger forces, as most cells started to move laterally in this situation (Supplementary Video 4). Nevertheless, these experiments confirmed that the pressure inside the aggregate rose in the LCC condition, and that this rise resulted in the cells generating a pushing force on the environment. This further supports the hypothesis that during the outbursts the cells are pushed by the pressure within aggregates, which is induced by the cell swelling.

To further test this hypothesis, we decided to increase the pressure acting on the aggregate^25^, thus reducing the pressure difference, which should then abolish the outbursts. This was achieved by adding 100 mg ml^*−*1^ Dextran 2 MDa to the outside, which generates a compressive force of about 18 kPa^26^, thereby exceeding the average pressure increase we measured using the elastic beads. Consistent with the pressure-driven outgrowth hypothesis, the addition of such an external pressure even in the LCC situation fully eradicated the outbursts within the same time scale (Figure 3a, Supplementary Figure 2). Interestingly, this counteracting surrounding pressure further reduced the cell motility and led to a sub-diffusive motion, which directly confirms that the cells were pressed together in a jamming-like fashion.

**Figure 3:**
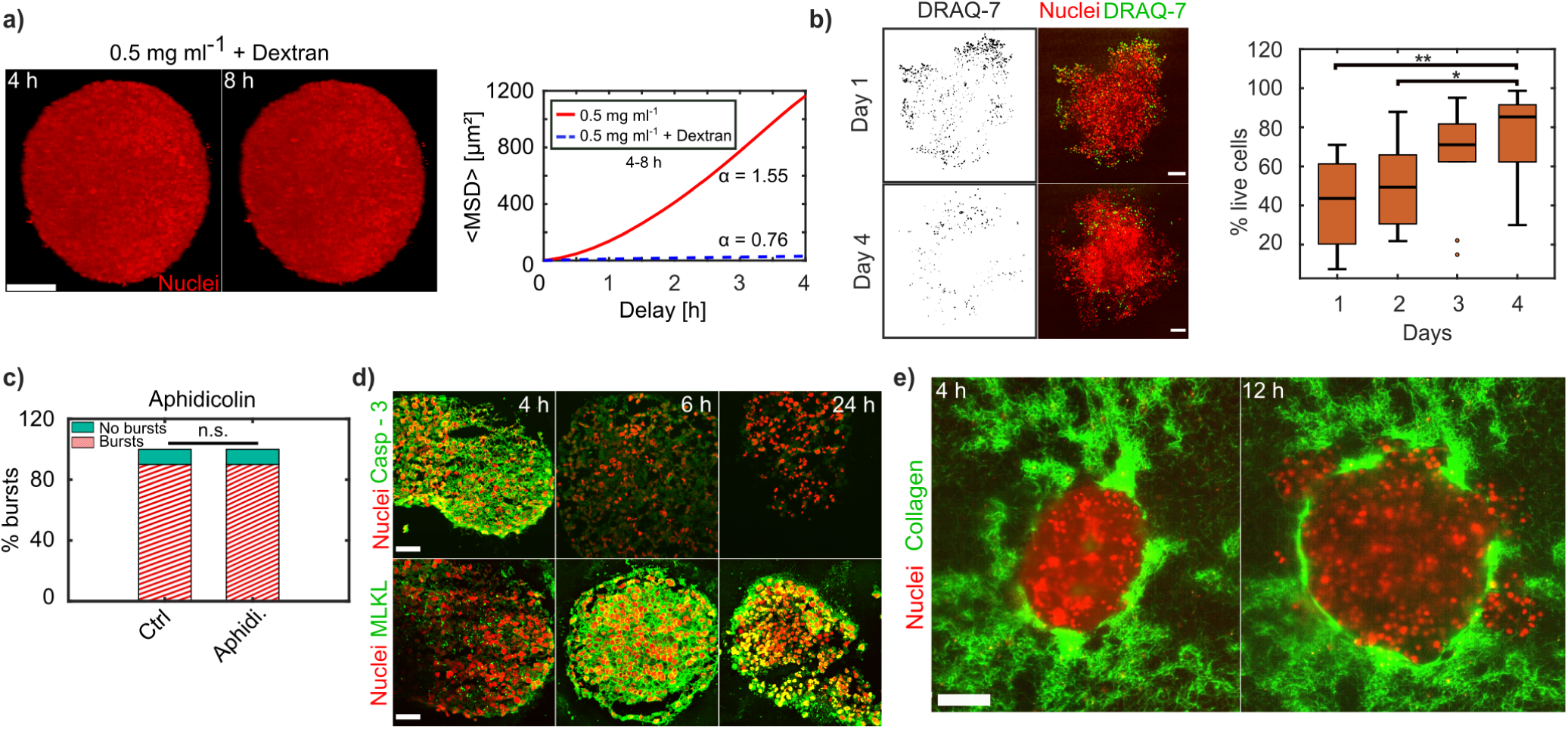
Collective pressure-driven invasion is independent of cell proliferation and correlates with cell death. **a)** Dextran pressure of 18 kPa in LCC stops bursts from aggregates, scale bar = 100 *µ*m. (MSD) between 4-8 h displaying sub-diffusive behaviour in LCC + dextran (N = 1) and super-diffusion in LCC (N = 10). **b)** HeLa H2B-mCherry aggregates stained for DRAQ-7 (green) on days 1 and 4 (left) post embedding. % cells alive from days 1 to 4 (N = 10 each). Medians indicated, scale bars = 100 *µ*m. **c)** % bursts in control (Ctrl) and with aphidicolin (Aphidi.) 5 *µ*g ml^*−*1^, N = 10 each. **d)** 12 *µ*m sections of aggregates stained for Caspase-3 and MLKL antibodies at 4, 6, 24 h, scale bars = 50 *µ*m. **e)** Collagen pockets seen at 4 and 12 hpe, scale bar = 100 *µ*m.

## Collective pressure-driven invasion is independent of cell proliferation and correlates with cell death

Although the pressure-driven effect provides an excellent explanation for the observed collective cell bursts, it does not illuminate the underlying mechanism. As proliferation might explain the volume increase, we arrested cells in S-phase of the cell cycle, by adding (5 *µ*g ml^*−*1^) aphidicolin. However, this did not reduce the burst phenotypes (Figure 3c), suggesting that proliferation is not required for the collective bursts. Another mechanism could be that in soft environments, like the LCC condition, cell death occurs via failed mechanotransduction^27, 28^. To test whether the cells within the aggregates underwent massive cell death we used the membrane-permeable dead cell stain DRAQ-7, which marks nuclei in the case of plasma membrane rupture. Indeed, we found that one day post embedding, a high number of cell nuclei stained positive for DRAQ-7 (Figure 3b), demonstrating increased cell death. Furthermore, cell death increased in the regions of the outbursts, suggesting a relation between bursts and reduced cell survival. Since DRAQ-7 is a classical necrosis marker, we further wanted to confirm whether apoptosis possibly preceded the cell death, as reported previously^29^. Staining against the apoptosis marker Caspase-3 was highest at 4 hpe as compared to mixed lineage kinase domain-like pseudokinase (MLKL, for necrosis) which drastically increased between 6-24 hpe (Figure 3d). Thus, these markers establish the series of events leading to cell death during the burst phase. Interestingly, when inspecting the live aggregates 4 days after bursts, the DRAQ-7 staining was largely absent, indicating that the overall cell viability had recovered, even though the environment remained the same. We quantified this recovery by recording the percentage of live cells as a function of days (Figure 3b, Supplementary Figure 4) and confirmed that already 2 days post embedding the viability increased and recovered to almost 90% living cells after 4 days.

As the current picture of collective migration is rooted in force transmission between cells by pulling forces, which is the opposite of the pushing force (pressure) we identified here, we wondered whether we would still observe the outbursts if we blocked cell-cell adhesion by a commonly used cadherin antibody (Supplementary Figure 2). Indeed, we observed that the bursts were even more pronounced when reducing the cell-cell adhesion, suggesting that the pressure-based collective migration mechanism is fundamentally different from classical collective migration schemes.

## Outburst regions are controlled by a limited mechanical resistance due to collagen in-homogeneities

We demonstrated that the bursts are pressure-driven, independent of cell proliferation or cell-cell adhesion, and generated by cell swelling that is triggered by the LCC. However, the reason for localised outbursts instead of a homogeneous aggregate expansion was still unclear. Motivated by the sandwich experiments, where we saw lateral motion of the cells while the aggregate was pushing in the PAA substrate, we hypothesised that in-homogeneities in the LCC structure could facilitate an expansion into regions with the least mechanical resistance. Indeed, when imaging the collagen after embedding the aggregate we saw large spatial concentration variability (Figure 3e) potentially due to the low overall collagen concentration, but this could have been also an outcome of the embedding process. Consistent with the hypothesis of bursts into regions of least resistance, we could precisely predict the bursts to occur in the directions of reduced collagen presence. To test whether simple swelling combined with the in-homogeneity of the collagen could fully explain the observed outbursts, we simulated the situation by modelling the aggregate as a cluster of spheres embedded in an in-homogeneous environment, mimicking the experimental situation (Figure 4a). When applying the observed volume increase to the individual cells as observed in the experiments (Supplementary Figure 3, 5), the simulations correctly reproduced the collective outbursts. This demonstrates that the model proposed here (Figure 4b) can explain the observations, even when only using the highly simplified situation of interacting spheres.

**Figure 4:**
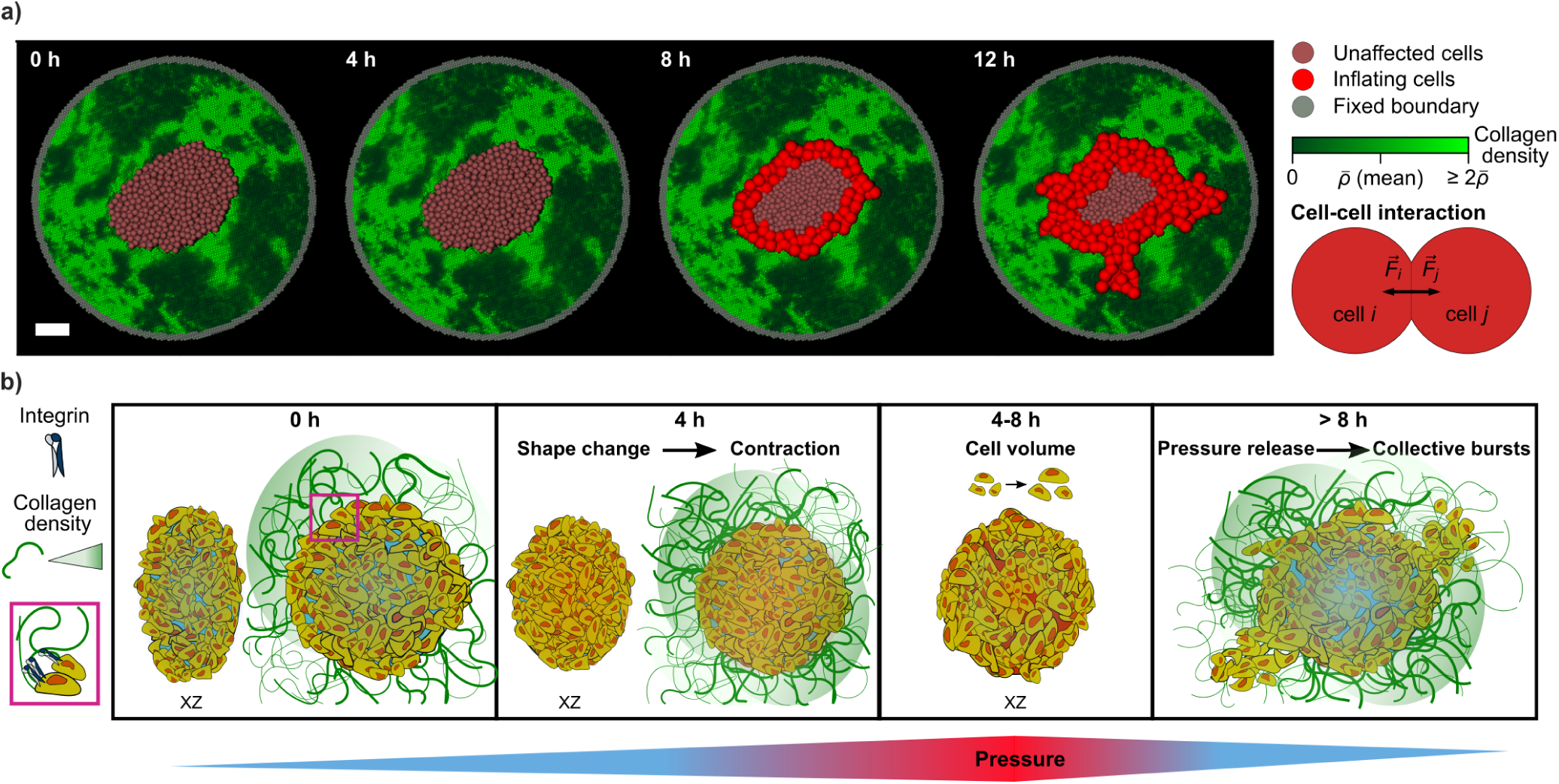
Bursts in simulation model, considering collagen in-homogeneity and cell swelling in LCC. **a)** Simulation results for the time evolution of a cancer spheroid. The initial distribution of cancer cells and collagen was chosen analogous to the experimental data (scale bar = 100 *µ*m). **b)** Sketch representing series of events over 0-8 h triggering collective pressure-driven burst-like invasion. **0 h** Cells at the periphery of oblate aggregates attach to the LCC via integrins (see legend and zoomed inset). Internal pressure remains low. **4 h** The ECM attachment sculpts a shape change. Pressure starts to be disturbed. **4-8 h** Volume of cells increases within aggregates raising the internal pressure. From **8 h** onwards, aggregates are unable to sustain the pressure increase and expel cells collectively in the directions of least mechanical resistance (green gradient (legend) depicts collagen density (in-homogeneity in LCC)).

## Conclusion

Here, we report a new, pressure-driven collective migration mechanism that is initiated by a contraction of cells in a low collagen concentration environment (Figure 4b). The contraction explains the observed initial collective and super-diffusive motion while the aggregate changed from an oblate to a more spherical shape. In this process, we suspect that the integrins failed to mechanically engage because of the LCC condition’s low mechanical resistance. Similar to anoikis^30^, such a failed engagement leads to massive cell death that is accompanied by cell swelling, which autonomously starts to increase the pressure inside the aggregate and pushes the cells into the environment in the direction of the least mechanical resistance. This model can explain all the observations and suggests a new mechanism of collective migration that is driven by pressure and, hence, is independent of cell-cell adhesion. In fact, this mechanism contradicts the standard paradigm for collective migration, but it is nonetheless a simple and almost intuitive outcome of pressure release. Although here we report a special situation in low collagen concentration matrices, the general pressure-driven burst mechanism might be highly relevant in the context of cancer cell invasion. Primary tumours are often encapsulated by a basement membrane that constraints the growing tumour. Upon increased proliferation, the pressure inside the tumour is known to increase. The basement membrane may rupture either by simple mechanical tension or due to active degradation by the cancer cells. In this situation the pressure might push the cells out in a similar way as demonstrated here, via cell outbursts, potentially producing drastic consequences while promoting metastasis.

## Methods

### Cell culture and aggregate preparation

HeLa cervical cancer cell lines stably transduced to express H2B-mCherry, H2B-mCherry and Life-Act GFP, H2B-RFP and *α*-tubulin (kind gift from Matthieu Piel), or H2B-mCherry and MyrPalm-GFP, via lentiviral transduction, were cultured using high glucose DMEM (Capricorn) medium supplemented with 10% (v/v) fetal bovine serum (FBS, Sigma-Aldrich) and 1% (v/v) penicillin-streptomycin solution (Gibco). Cells were stored at 37 °C in a humidified atmosphere with 5% CO_2_, and split on reaching a confluency of *>* 60%. Split ratios were either 1:5, 1:10 or 1:20.

Cancer aggregates were prepared in a 48-well plate (Greiner Bio-one) as previously described^17^. Plates coated with 150 *µ*l of 1% ultra-pure agarose (Invitrogen) in each well were cooled for 30 min prior to adding 1 ml of cell suspension containing 2000-2500 cells. The aggregates were collected between days 2-4 and imaged.

### Collagen polymerisation

Collagen concentrations of either 0.5 mg ml^*−*1^ or 2.5 mg ml^*−*1^ were prepared using a mixture of 10x phosphate-buffered saline (PBS, Sigma-Aldrich) (diluted to 1x in the final volume prepared), rat tail collagen I (Corning, stock: 3.77 mg ml^*−*1^), and cell culture medium. The polymerisation was activated by adding 1 M NaOH to attain a pH of 7.5. All solutions used were stored in 4 °C before, and kept on ice under sterile conditions while preparing the polymerisation mix. CO_2_ independent medium supplemented with 10% (v/v) fetal bovine serum (FBS, Sigma-Aldrich) and 1% (v/v) penicillin-streptomycin solution (Gibco) was used instead of DMEM for experiments performed with either light sheet or spinning disk confocal microscopes. In order to mark the collagen fibres, a collagen-binding adhesion protein 35-Oregon Green 488 (CNA35-OG488, stock: 8.2 mg ml^*−*1^, a kind gift from Gijsje Koenderink) was used at 1:20 of the final collagen concentration for either 0.5 or 2.5 mg ml^*−*1^ as required.

### Aggregate sample preparation and multi-view imaging

To image the aggregate at sub-cellular resolution we used a dual illumination light sheet microscope (Zeiss Z.1, 20x objective, N.A. 1.0). FEP (Fluorinated Ethylene Propylene) capillaries (Proliquid, dimensions: 1.6 mm *×* 2.4 mm, width: 0.4 mm) along with Teflon plungers (Zeiss, size 3) suitable for incorporating into the microscope’s sample holder were used. Once the cells aggregated on day 2 or 3, they were first removed from the 48-well plate and transferred to a Petri dish (92 mm *×* 16 mm, Sarstedt). 100 *µ*l of collagen polymerisation mixture was introduced into the Petri dish at room temperature (22-24 °C). Aggregates were then transferred into this collagen droplet with minimal medium to avoid dilution of the droplet. Finally, the aggregates were pulled into the FEP capillary via the suction force provided by the plunger. The capillary was then repeatedly rotated horizontally by hand, holding the part where the aggregate suspends in the polymerising collagen until it was stably centred (15-20 min) within collagen fibres. 1% ultra-pure agarose (Invitrogen) was then plugged at the bottom to avoid collagen to seep out of the capillaries over time. The sample capillaries were then introduced into the sample chamber of the light sheet microscope filled with CO_2_ independent medium supplemented with 10% (v/v) fetal bovine serum (FBS, Sigma-Aldrich) and 1% (v/v) penicillin-streptomycin solution (Gibco) equilibrated to 37 °C, 30 min prior to imaging.

Nuclei of cells forming aggregates within polymerised collagen with concentration of 0.5 mg ml^*−*1^ (LCC) were imaged with a 15 min time interval and those of 2.5 mg ml^*−*1^ (HCC) were imaged with both 15 min (N = 5) and 1 h (N = 5) time intervals. Multi-views of the samples were acquired at 4 different angles (multiples of 90°). Each view comprised z-stacks in the range of 600-800 *µ*m, separated by an optimal distance of 2 *µ*m. The lateral resolution for image stacks was in the range 0.45-0.56 *µ*m and the light sheet thickness was *≈*6.47 *µ*m. Fluorescent beads (505 nm, Invitrogen; 561 nm, Micromod) of sub-cellular resolution were embedded in the collagen in order to register the different views. Registration was done using the ‘Multiview Reconstruction’^31, 32^ plug-in from Fiji^33^. The point spread function of the microscope could be estimated from these registration beads using the plug-in, and images were deconvolved and downsampled 2x or 3x for further analysis.

### Single cells sample preparation and imaging

The preparation was done immediately after passaging HeLa cells (stably expressing H2B-mCherry and LifeAct-GFP) using culture media. Cell pellets were re-suspended in 1 ml CO_2_ independent medium. LCC polymerisation mixture volume of 100 *µ*l consisting of 80,000 cells was prepared by adjusting the volume of the medium (for collagen) with respect to the volume of cells added in the mixture. The sample mix was not vortexed to prevent any damage to cells. Once the pH was adjusted to 7.5, the sample mix was pulled into the FEP capillary and rotated until the collagen polymerised (similar to aggregate preparation) and imaged with the light sheet microscope.

### Cell detection and tracking in aggregates

Nuclei marked for H2B-mCherry or H2B-RFP were tracked via ‘Autoregressive Motion’ algorithm in Imaris 9.3.0 through the ‘Spot detection and tracking’ method. XY diameters were estimated from the ‘Slice’ view and thresholds were set for the minimum intensity and quality of detection in order to filter for noise spots detected. All further analyses of tracks were done using MATLAB R2016a. Nuclei that were unable to be segmented since their intensities did not match the threshold criteria, were omitted, and track positions were interpolated for missed links (typically 1-2).

### Track analysis

#### Absolute velocity

The absolute velocity 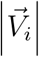 for every particle *i* in 3D (x, y, z) was calculated as 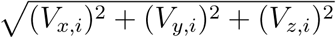 over time. Averages and s.e.m. were then acquired for every time point.

#### Mean squared displacement (MSD)

The MSD of all particles over the time delays was calculated using a previously described method^34^. The time window based MSD was an adapted version of the method, where it was only applied on the tracks within the respective time windows, given a window size of 10 time points (total 49 time points (12 h) with 15 min time interval). We fit a linear function *f(x)* = *α x* + 6*D* to the logarithmic data of the equation ⟨x(*τ*)^2^⟩ = 6*Dτ*^*α*^ in the moving time window (2.5 h) to attain the exponent *α* and the diffusivity constant *D* as a function of *τ* (delays).

#### Voronoi volume of cells within aggregates

Based on tracked nuclei positions, a MATLAB function ‘N-D Voronoi diagram’ was applied to retrieve an estimate for the cell borders. Next, all cell borders per time point were fit by an ellipsoid (ellipsoid fit, https://www.mathworks.com/matlabcentral/fileexchange/24693-ellipsoid-fit,MATLAB Central File Exchange) to extract an approximation for the cell volume.

#### Velocity correlation

The velocity vectors of all particles *i* per time point *t* were correlated with every other particle *j* and normalised by the square of the mean absolute velocities of *i* and *j*, leading to the correlation function

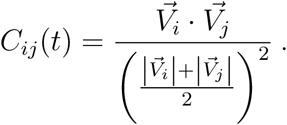

*C*_*ij*_(*t*) was binned over the distances between particles *i, j* (bin size = 20 *µ*m) and the averages per bin were acquired for each time point. Next, ⟨*C*_*ij*_(*t*)⟩ up to a binned distance of 150 *µ*m was fit by an exponential function *f(x)* = *a e*^*−b x*^, where the correlation length is defined as 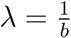.

### Spherical harmonics for shape change and aggregate volume

Spherical harmonics are a complete set of orthogonal functions defined on the unit sphere and are extracted using the Python toolbox shtools^35^. To determine the shape and volume changes of the aggregates over time, their surfaces were described using the expansion

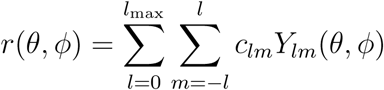

where *r* denotes the distance of a point on the surface from the center of mass of the aggregate, *θ* and *φ* are the spherical coordinate angles, *c*_*lm*_ the spherical harmonic coefficients, *Y*_*lm*_ the spherical harmonics, *l* and *m* their degree and order, respectively, and *l*_max_ the maximum degree of the expansion.

The enclosed volumes of these surfaces were determined by the integral

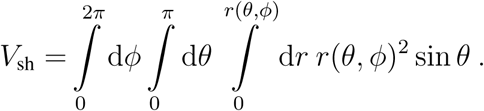

The higher the limit of the sum *l*_max_, the finer the details of a surface that can be described by the spherical harmonics. Due to this reason, on the raw data, a variable expansion degree was used as the shape of the aggregates evolved, starting with typically *l*_max_ = 4 for the oblate shape at earlier time points and reaching up to *l*_max_ = 15 for the intricate shapes occurring at later time points as a consequence of the bursts.

We observed that such a variation in the spherical harmonic expansion degrees on raw data over time gave rise to offsets/artefacts in the volume detection, that had to be corrected. Assuming that drastic changes in volume do not occur at short time scales, and understanding the error source due to the expansion degree changes, we readjusted the volume by adding the difference caused by the offset to the consecutive time points. To verify if the detected volumes were influenced by the raw experimental data, we used a fixed expansion degree of *l*_max_ = 15 and determined the volumes based on aggregate surfaces estimated from tracked nuclei positions. The results were in close agreement with those from the raw data.

### Volume measurement of single cells

To measure the volume of HeLa single cells in LCC, stably expressing H2B-mCherry and LifeAct-GFP were used. The cell shapes via LifeAct-GFP were 3D segmented using CellProfiler 3.1.9^36^ on downsampled data (2.78 *µ*m pixel^*−*1^). A global ‘Otsu’ threshold was applied to segregate pixel intensities to background or foreground. This was followed by a standard ‘Watershed’ algorithm and segmented objects were measured for shape volume. Exported measurements were extracted and represented using MATLAB R2016a.

### Elastic beads as pressure sensors

A custom made^37^ water-in-oil emulsion was used to prepare the inert PAA beads of Young’s modulus *E* = 3.9 *±* 0.9 kPa, by injecting the water-based bead mix in oil dissolved in n-Hexane (Supelco). The PAA beads were fluorescently labelled (Atto 488 NHS-Ester, Atto-Tec) several times until sufficient fluorescent signal was achieved. HeLa H2B-mCherry aggregates were prepared (Methods) such that each well of the 48-well plate comprised *≈*30 PAA beads. Aggregates with randomly incorporated elastic beads were imaged in the same way as mentioned before (Methods).

To calculate the pressure applied on the elastic beads by the cells within aggregates, we retrieved first the main force dipole deforming the bead surfaces using spherical harmonics expansions. For this, it is sufficient to use the spherical harmonic components *Y*_00_ and *Y*_20_ as

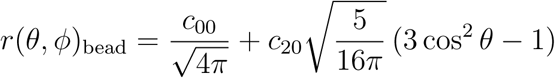

where the first term corresponds to the radius of the beads by 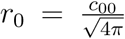 and the second term describes the major uni-axial deformation of a sphere. The surfaces of the beads were expanded in the 0^*th*^ and 2^*nd*^ degrees of the spherical harmonics using the above function, and the surfaces were rotated in such a way that the *c*_20_ coefficient was minimized, aligning then the main axis of compression with the z-axis. Using this expression as an ansatz to tackle the elastic problem on a sphere as expressed in the Navier-Cauchy equation, we arrived at the force dipole acting on the bead

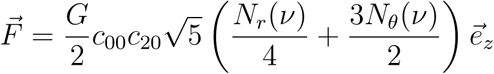

With 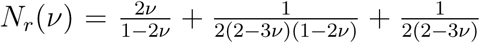 and 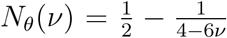, and ν and *G* being Poisson ratio and shear modulus of the PAA beads, respectively.

The pressure was then estimated from the relation 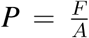 with 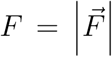 and *A* is taken as the surface of the bead,

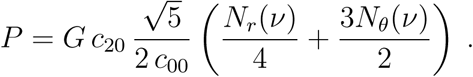

### Measuring the internal pressure by confining aggregates

HeLa H2B-mCherry day 2 aggregates were equilibrated for 1 h in CO_2_ independent medium containing 10% (v/v) fetal bovine serum (FBS, Sigma-Aldrich) and 1% (v/v) penicillin-streptomycin solution (Gibco) at 37 °C prior to using them for sandwich experiments. The sandwich consists of a bottom layer of PAA gel (*E ≈* 1800 Pa) and a top layer of a 12 mm round glass cover slip (VWR).

#### Bottom layer

A 35 mm glass-bottom dish (Greiner Bio-one) was first thoroughly cleaned with 0.1 M NaOH, silanized and washed twice. The dishes were then treated with 25% glutaraldehyde for 30 min. A pre-gel mixture of 40% acrylamide and 2% bis-acrylamide in a 2:1 ratio, along with 4 *µ*l of 99% acrylic acid was prepared and diluted with 65% PBS (Sigma-Aldrich) to achieve the required Young’s modulus (*E ≈*1800 Pa). 10 *µ*l fluorescent beads (505 nm, Invitrogen) were added to the pre-gel mix for traction force microscopy^22, 23^. The free radical polymerisation was triggered by adding 5 *µ*l ammonium persulfate and 1.5 *µ*l tetramethylethylenediamine. A 5 *µ*l drop of the final pre-gel mix was added to the centre of the glass-bottom dish and spread evenly by placing a 12 mm round cover slip on top for 20 min. The cover slip was then slid away from the gel after adding 65% PBS to the dish. For all experiments the PBS was aspirated completely and the gel was air-dried for 10 min before use.

#### Top layer

The top layer of the confinement was coated overnight with 50 *µ*g ml^*−*1^ collagen I (Corning, stock: 4.88 mg ml^*−*1^) in 0.2 M acetic acid containing fluorescent beads (*≈*1%). 100 *µ*l of the collagen solution prepared was then added to a Petri dish (92 *×* 16 mm Sarstedt), to which 2-3 aggregates were introduced, avoiding dilution with medium. The aggregates were then transferred one by one with *≈*20 *µ*l collagen volume to the PAA gel.

#### Confinement and Imaging

Images of the aggregates were taken using a spinning disk confocal system (CSU-W1 Yokogawa, Intelligent Imaging Innovations Inc.) via the Slidebook 6 software (3i) equipped with an inverted microscope (Nikon Eclipse Ti-E) and a CMOS camera (Orca-flash4.0v2, Hamamatsu Photonics K.K.). The microscope comprises a custom-built heating chamber, which was maintained at 37 °C during imaging. In order to confine the aggregates within a sandwich-like model, the top layer was adhered with the help of korasilon paste (Kurt Obermeier GmbH) to a custom-built device. The glass-bottom dish containing aggregates embedded in PAA gel was placed inside the device, parallel to the top. The device was then introduced within the heating chamber of the microscope.

The top layer was moved down until the upper periphery of the aggregates, such that they were only confined and not compressed. Once the positions of the aggregates were stabilised, 3 ml of CO_2_ independent medium was added. z-stacks with intervals of 0.5 or 1 *µ*m were acquired from the bottom to the top layer, with a time interval of 30 min for *≈*8 h. For control experiments, no collagen was involved for the bottom or the top layers.

### Contractility measurements

To obtain the forces that the spheroid exerts on its environment, we adapted a recently published procedure^20^ to the spheroid invasion system. Briefly, fluorescently labelled particles (100 nm diameter) embedded in the collagen matrix were tracked over time to reconstruct the local deformation of the hydrogel. Using macro-rheology, the non-linear response of the collagen gel was measured and used to create a lookup table that relates the observed deformation to the exerted pressure. Averaging over all directions, a mean contractility (pressure *×* surface area, *µN*) and s.e.m. were obtained.

### Applying external pressure in LCC

0.5 mg ml^*−*1^ collagen was polymerised along with 100 mg ml^*−*1^ Dextran 2 MDa (Dextran T2000, Pharmacosmos) to achieve a pressure of *≈*18 kPa^26^. Aggregates were then subject to this environment and imaged for 12 h using the light sheet microscope (Zeiss Z.1, 20x objective, N.A. 1.0).

### Chemical reagents to inhibit aggregate bursts

Aphidicolin (Sigma-Aldrich, A4487) and E-cadherin Antibody (G-10, SCBT) were used at working concentrations of 5 *µ*g ml^*−*1^ and 10 *µ*g ml^*−*1^, respectively. Aggregates to be treated with aphidicolin were pre-incubated with the drug along with culture media for 1 h at 37 °C in a humidified atmosphere with 5% CO_2_. After an hour, aphidicolin pre-incubated aggregates and E-cadherin untreated aggregates were embedded in FEP capillaries containing LCC along with the respective drug/chemical (N = 10 each). Sample capillaries were immersed in 1 ml culture medium (with or without drug/chemical (control)) and stored at 37 °C in a humidified atmosphere with 5% CO_2_ overnight. After 24 h, samples were slid out of the capillaries carefully into a Petri dish and bright-field images were taken using an inverted microscope (Vert.A1 Zeiss, Axio).

### Quantifying cell survival

FEP capillaries consisting of HeLa (stably expressing H2B-mCherry, or H2B-mCherry and LifeAct-GFP) aggregates that had undergone a burst-like invasion within 8 h of sample preparation (Methods), were continued to be stored at 37 °C in a humidified atmosphere with 5% CO_2_ from days 1-4 (N = 10, for each day). After each day, the capillaries were removed and samples were pushed out carefully into a Petri dish (60/15 mm, Greiner Bio-one), after which 100 *µ*l of culture medium was added on top of each of the aggregates (still in LCC) to prevent drying out. From here on the samples were protected from light, and DRAQ-7 (Thermofisher) was administered at 1:100 ratio to every sample droplet and the Petri dish was stored at room temperature for 10 min. Spinning disk confocal images of the nuclei with lasers 561 nm and 647 nm (DRAQ-7) were acquired with the Slidebook 6 software (3i) using an inverted microscope (Nikon Eclipse Ti-E) equipped with a CSU-W1 spinning disk head (Yokogawa) and a scientific CMOS camera (Prime BSI, Photometrics).

Nuclei were detected using Laplacian of Gaussian detection algorithm from Trackmate^38^. The number of dead cells was subtracted from the total number of cells detected, to retrieve the % live cells.

### Cryosectioning and immunostaining aggregates

Capillaries consisting of HeLa H2B-mCherry day 3 aggregates in 0.5 mg ml^*−*1^ collagen were removed from the medium at 4, 6 and 24 h and immersed in 1x PBS (3 times) to wash. The capillaries were dipped into 4% Paraformaldehyde 1 ml for 30 min. Next, capillaries were washed again 3 times with 1x PBS. To preserve the structure of the bursts in collagen, the capillaries were subjected to 500 *µ*l optimal cutting temperature mounting medium (Sakura) for 1 h. They were then partially snap-frozen with liquid nitrogen vapour. After 30 s at room temperature, the frozen samples were pushed out of the capillaries into a block of liquid optimal cutting temperature mounting medium and centred before it was snap-frozen completely. Samples were stored at −80 °C until further processing.

Frozen aggregates were sectioned, mounted as 12 *µ*m sections and re-hydrated. The samples were blocked for 1 h at room temperature using 1x PBS supplemented with 10% goat serum (Sigma) and 0.2% Triton-X-100 (Carl Roth). Subsequently, the samples were incubated with the primary antibody (monoclonal mouse anti-Caspase-3, 1:100, Santa Cruz, sc-56053; monoclonal mouse anti-MLKL, 1:100, Proteintech, 66675-1-Ig) diluted in blocking solution over night at 4 °C. After three washes with PBS, samples were incubated with the secondary antibody (polyclonal goat anti-mouse IgG1, 1:500, ThermoFisher) diluted in blocking solution for 45 min at room temperature. Samples were finally washed with PBS three times and confocal images were acquired with Slide-book 6 software (3i) using an inverted microscope (Nikon Eclipse Ti-E) equipped with a CSU-W1 spinning disk head (Yokogawa) and a scientific CMOS camera (Prime BSI, Photometrics).

### Statistics

Data noise emerging as part of the image acquisition over a 12 h time series was reduced by applying a median smoothing (median of every 5 consecutive time points). The application of this smoothing function is mentioned in each figure legend. All error shades (s.e.m.) in plots were generated using an adapted function (shaded area error bar plot, https://www.mathworks.com/matlabcentral/fileexchange/58262-shaded-area-error-bar-plot, MAT-LAB Central File Exchange).

A chi-squared test was performed to compare between the categories ‘drug/chemical’ (N = 10) and ‘control’ (N = 10). All other data were tested for significance using the two-sample independent t-test, considering significance of p*≤*0.05. The significant p-values were categorised as p*>*0.05 = ‘n.s.’, p*≤*0.05 = ‘*’, p*≤*0.01 = ‘**’, p*≤*0.0001 = ‘***’, and p*≤*0.00001 = ‘****’.

## Data availability

All relevant data supporting the findings of this study are available from the corresponding author upon reasonable request.

## Code availability

The codes that were used to analyse the data or to perform the simulations are available from the corresponding author upon reasonable request.

## Acknowledgements

We are very thankful to Matthieu Piel and Roland Wedlich-Söldner for offering us HeLa H2B-RFP *α*-tubulin-GFP and HeLa wildtype cell lines, respectively. We are grateful to Gijsje Koenderink for providing us with CNA35-OG488 to mark collagen fibres successfully. We thank Erez Raz, Johanna Ivaska and Peter Friedl for helpful discussions. R.W. and T.B. are funded by the Deutsche Forschungsgemeinschaft (DFG, German Research Foundation) – Grant No. WI 4170/3-1 (R.W.) and Project-ID 450595133 (T.B.). The simulations for this work were performed on the computer cluster PALMA II of the University of Münster. S.R. acknowledges funding by the fund Innovative Medical Research of the University of Münster Medical School, BE 12 16 09. This work was funded by the European Research Council ERC-Consolidator grant PolarizeMe (771201).

## Author contributions

S.R., A.S. and A.M. carried out experiments and analysed data. S.B. and R.W. performed the simulations and the comparative nuclei-based spherical harmonics analysis. Traction stresses and spherical harmonics expansions were extracted from the raw data by S.R. using custom-built programs by M.B. and A.J., respectively. The sandwich device was custom made by F.A. Sectioning and immunostainings were made by A.H. Non-linear collagen responses were measured by B.V. and S.R. T.B. designed and supervised the study. S.R., S.B., R.W. and T.B. wrote the manuscript.

