## Supplementary Information for "Pressure drives rapid burst-like collective migration from 3D cancer aggregates"

Swetha Raghuraman[1*], Ann-Sophie Schubert[1], Stephan Bröker[3], Alejandro Jurado[1,2], Annika Müller[1], Matthias Brandt[1], Bart E. Vos[1,2], Arne D. Hofemeier[1], Fatemeh Abbasi[1,2], Martin Stehling[4], Raphael Wittkowski[3] & Timo Betz[1,2]

[1]*Institute of Cell Biology, ZMBE, University of Münster, Münster, Germany*

[2]*3$^{rd}$ Institute of Physics, University of Göttingen, Göttingen, Germany*

[3]*Institute of Theoretical Physics, Center for Soft Nanoscience, University of Münster, Münster, Germany*

[4]*Max Planck Institute for Molecular Biomedicine, Münster, Germany*

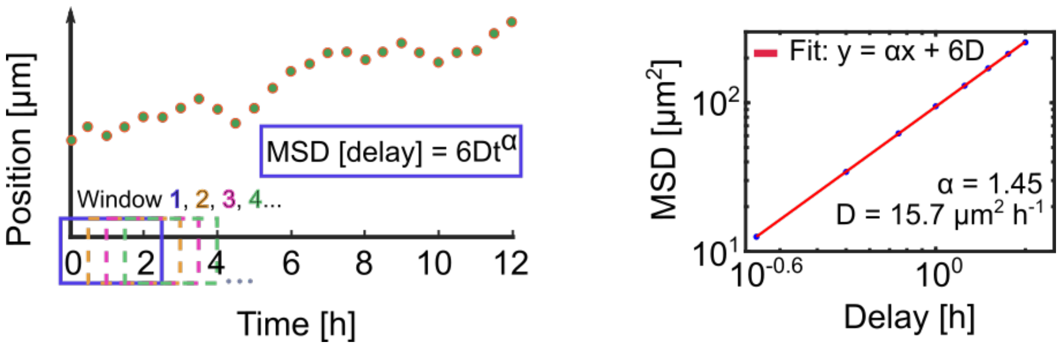


Supplementary Figure 1: **Time window MSD.** Tracks from a sliding time window of 2.5 h over 12 h are retrieved and MSD is calculated (left). The expression $x(\tau)^2$ = 6 D $\tau^{\alpha}$ is fit onto the log(MSD) to extract the exponent $\alpha$ and diffusion coefficient D (right). Shown here is the fit for the last time window of the series.

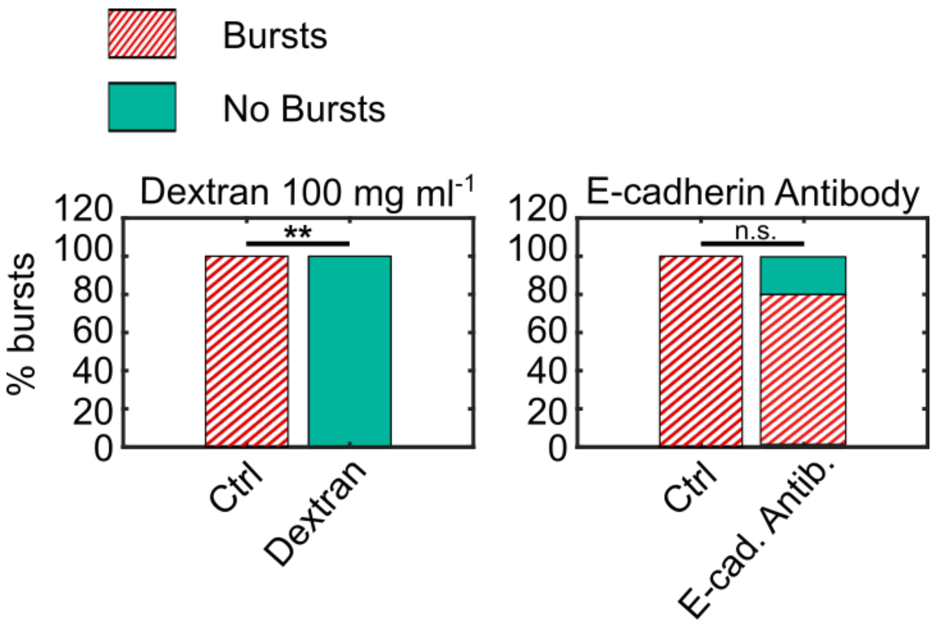


Supplementary Figure 2: **External pressure and not cell-cell adhesion blocking stops bursts.** An external pressure of 18 kPa using dextran (100 mg $ml^{-1}$) added in LCC stopped bursts completely (left). E-cadherin antibody (G-10) (80% bursts, right) when added did not arrest the bursts. Control (Ctrl) N = 3, Dextran/Antibody N = 10.

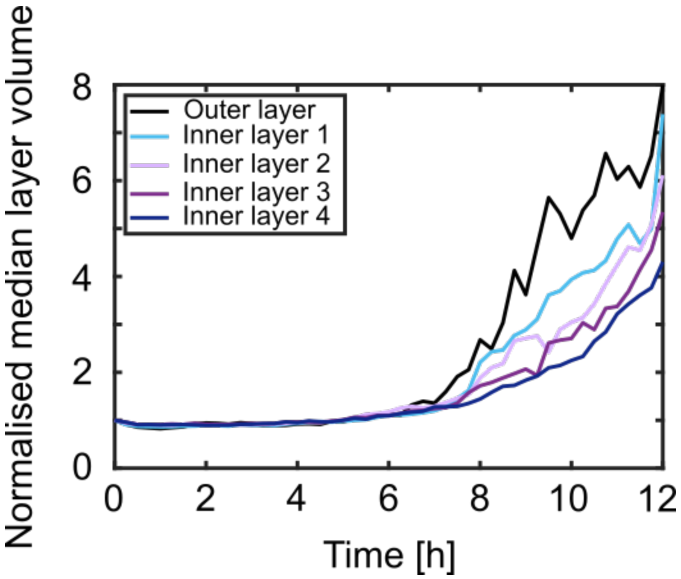


Supplementary Figure 3: **Aggregate volume in layers.** Voronoi-based median of volumes over time was retrieved for individual layers (up to 5, normalised to time 0) within aggregates starting from the outermost cells, N = 10.

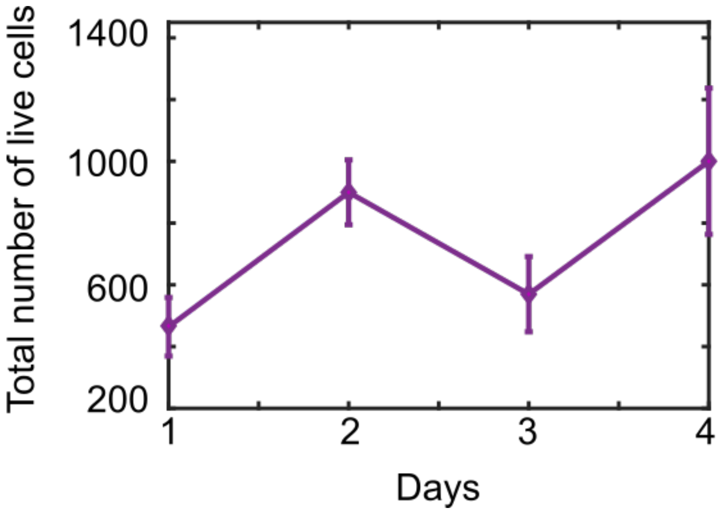


Supplementary Figure 4: **Number of live cells increases over days.** Total number of live cells over days post embedding of aggregates in LCC. N = 10, s.e.m.

### 1 Setup of the simulated system

In the computer simulations of the time-evolution of an initial cancer aggregate, the cancer cells are surrounded by collagen that is confined by a fixed spherical boundary at a large distance (see Supplementary Figure 5). As in the experiments, the aggregate has an initial shape of a slight oblate, with a size of $385\,\mu m$, $318\,\mu m$, and $271\,\mu m$ in the $x$, $y$, and $z$ direction, respectively. Furthermore, the surrounding sphere of collagen has a radius of $400\,\mu m$, its centre coincides with the centre of mass of the cancer aggregate, and its boundary has a thickness of $20\,\mu m$ (see Supplementary Figure 5).

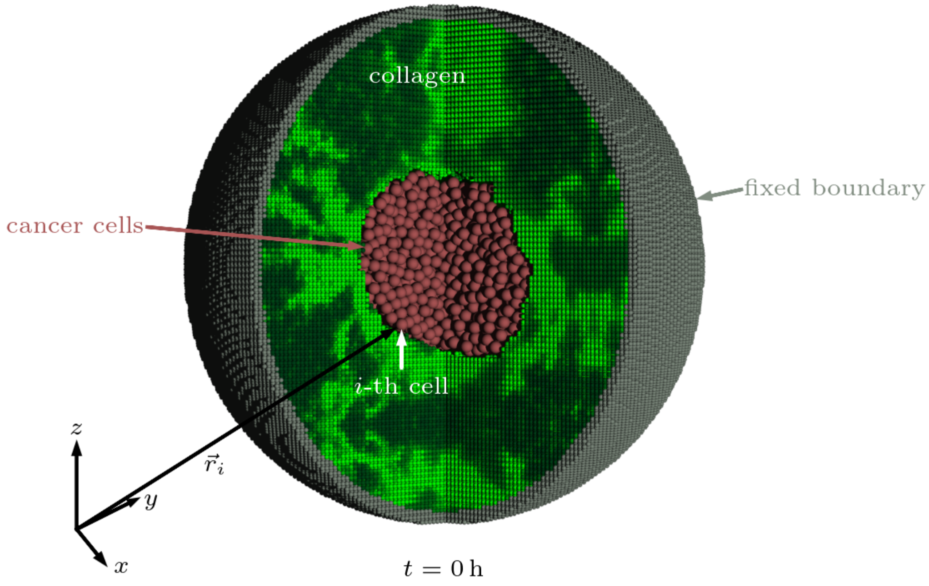


Supplementary Figure 5: **Sketch of the initial setup of the simulations.** The cells in the centre are surrounded by collagen and this is confined by a fixed spherical boundary.

**Cancer cells.** The cells are approximated as spherical particles and, as in the experiments, their radii $R(t)$ can increase as a function of time $t$. We choose their initial radii as $R_0 = R(0) = 10\,\mu\mathrm{m}$. To be close to a realistic situation, for the initial distribution of the cells, we place the spherical particles at the positions of the nuclei of the cells that are observed in the experiments at the initial point in time.

**Collagen.** For the collagen, we use a bead-spring model, i.e., a grid of small spherical particles that are connected with springs. The particles are initially placed on a body centred cubic lattice and each particle is connected with springs to its nearest and next-nearest neighbours. The local stiffness of the collagen is chosen according to the experiments at the start of the observation period, and remains unchanged during the simulations. For this purpose, images of the collagen from the beginning of the experiments are used and the local density of the collagen is extracted voxel by voxel by assuming that it is a linear function of the brightness of a voxel. The mean of the densities corresponding to a cube of 8 voxels in each dimension, whose centre is closest to a particle of the grid, is then assigned to this particle, and the spring constant of a spring connecting two particles is then chosen proportional to the mean of the local collagen densities assigned to these particles. Since the voxel size is $1.1165\,\mu\mathrm{m}$, we choose a lattice constant of $a = 8 \cdot 1.1165\,\mu\mathrm{m} = 8.932\,\mu\mathrm{m}$. The distance to the nearest neighbour of a grid particle is then $a_{\mathrm{nn}} = a\sqrt{3}/2$ and the distance to the next-nearest neighbour is $a_{\mathrm{nnn}} = a$.

For consistence with the experiments, the proportionality constants for the spring constants that correspond to nearest-neighbour and next-nearest-neighbour interactions, respectively, are chosen so that the storage modulus of the collagen grid is in line with the value $10\,\mathrm{Pa}$ measured in the experiments for shear of a half percent. To imitate this storage modulus in the simulations, the spring constant of the springs between nearest neighbours at positions $\vec{r}$ and $\vec{r}'$ is chosen as the function $k_{\mathrm{nn}}(\vec{r},\vec{r}') = 3000a((\rho_{\mathrm{loc}}(\vec{r}) + \rho_{\mathrm{loc}}(\vec{r}'))/(2\bar{\rho}_0))\,\mathrm{Pa}$ and the spring constant for the springs between next-nearest neighbours is chosen as the function $k_{\mathrm{nnn}}(\vec{r},\vec{r}') = k_{\mathrm{nn}}(\vec{r},\vec{r}')/2$ with the initial local collagen density $\rho_{\mathrm{loc}}(\vec{r})$ according to the experiments and the spatially averaged initial collagen density $\bar{\rho}_0 = 0.5\,\mathrm{g\,l^{-1}}$. As the experimental data correspond to a cuboidal domain with edge length $712\,\mathrm{\mu m}$, $807\,\mathrm{\mu m}$, and $580\,\mathrm{\mu m}$ in $x$, $y$, and $z$ direction, respectively, and thus do not cover the full simulation domain, the experimental data for the initial local collagen density are extrapolated by mirroring at the domain's faces.

To take the viscoelasticity of collagen into account, the springs break permanently if they are stretched for more than five percent. The interaction force for a grid particle $i$ interacting with other grid particles is therefore

$$\vec{F}_{\mathrm{grid},i} = \sum_{j\in\mathcal{I}_{\mathrm{nn}}(i)} \vec{f}_{\mathrm{nn},i,j} + \sum_{j\in\mathcal{I}_{\mathrm{nnn}}(i)} \vec{f}_{\mathrm{nnn},i,j} \tag{1}$$

with the sets $\mathcal{I}_{\mathrm{nn}}(i)$ and $\mathcal{I}_{\mathrm{nnn}}(i)$ of the indices of the nearest neighbours and next-nearest neighbours of the $i$-th particle, respectively. Here, the particle-particle interaction forces are given by

$$\vec{f}_{\mathrm{nn},i,j} = \frac{\vec{r}_j - \vec{r}_i}{\|\vec{r}_j - \vec{r}_i\|}(\|\vec{r}_j - \vec{r}_i\| - a_{\mathrm{nn}})k_{\mathrm{nn}}(\vec{r}_i,\vec{r}_j)\Theta(1.05a_{\mathrm{nn}} - \|\vec{r}_j - \vec{r}_i\|), \tag{2}$$

$$\vec{f}_{\mathrm{nnn},i,j} = \frac{\vec{r}_j - \vec{r}_i}{\|\vec{r}_j - \vec{r}_i\|}(\|\vec{r}_j - \vec{r}_i\| - a_{\mathrm{nnn}})k_{\mathrm{nnn}}(\vec{r}_i,\vec{r}_j)\Theta(1.05a_{\mathrm{nnn}} - \|\vec{r}_j - \vec{r}_i\|) \tag{3}$$

with the position of the $i$-th particle $\vec{r}_i$, the position of the $j$-th particle $\vec{r}_j$, and the Heaviside function $\Theta$.

**Boundary.** The fixed boundary confines the system. It is incorporated into the simulations by particles that are similar to those from the collagen grid and continue the initial collagen grid in all directions, but their positions are fixed to avoid motion of the boundary particles.

### 2 Equations of motion

We describe the motion of the cells and particles of the collagen grid as overdamped dynamics. Brownian motion is neglected, since one can expect it to have no significant effect on the cells' motion. We confirmed the correctness of this assumption by additional simulations that take Brownian motion into account.

The equations of motion of the cells are given by

$$\dot{\vec{r}}_i = \frac{1}{\gamma_{\mathrm{cell}}} \sum_{j \neq i} \vec{F}_{\mathrm{int},i,j} \tag{4}$$

with the position of the $i$-th cell $\vec{r}_i$, the damping coefficient $\gamma_{\mathrm{cell}} = 100\,\mathrm{nN\,\mu m^{-1}\,s}$, the sum over the other cells and collagen-grid particles $\sum_{j \neq i}$, and the force of interaction with another cell or grid particle $\vec{F}_{\mathrm{int},i,j}$. An overdot denotes a derivation with respect to time. The interaction force of the $i$-th cell interacting with another cell or grid particle with index $j$ is chosen as

$$\vec{F}_{\mathrm{int},i,j} = \begin{cases} \frac{\vec{r}_i - \vec{r}_j}{d_{ij}} \frac{R_i + R_j - d_{ij}}{R_0} \epsilon_{\mathrm{rep}}, & \text{if } d_{ij} \leq R_i + R_j, \\ -\frac{\vec{r}_i - \vec{r}_j}{d_{ij}} \epsilon_{\mathrm{att}}, & \text{if } R_i + R_j < d_{ij} < 1.05(R_i + R_j) \\ & \text{and } R_i = R_j = R_0, \\ 0, & \text{else} \end{cases} \tag{5}$$

with the centre-to-centre distance $d_{ij} = \|\vec{r}_i - \vec{r}_j\|$ of particles $i$ and $j$, their radii $R_i$ and $R_j$, respectively, the repulsive-interaction strength $\epsilon_{\mathrm{rep}} = 50\,\mathrm{nN}$[1], and the attractive-interaction strength $\epsilon_{\mathrm{att}} = 0.2\,\mathrm{nN}$[2]. Equation (5) describes a repulsive interaction for small centre-to-centre distances and an attractive interaction when the distance is slightly larger than the sum of the particles' interaction radii. The attractive interaction is omitted for inflating cells (see section 3). While the interaction radius of a cell equals its current geometric radius, the interaction radius of a grid particle is always $R_0$.

The equations of motion of the grid particles are given by

$$\dot{\vec{r}}_i = \frac{1}{\gamma_{\mathrm{grid}}}\left(\vec{F}_{\mathrm{grid},i} + \sum_{j=1}^{N_{\mathrm{c}}} \vec{F}_{\mathrm{int},i,j}\right) \tag{6}$$

with the damping coefficient for grid particles $\gamma_{\mathrm{grid}} = \gamma_{\mathrm{cell}}$ and the total number of cells $N_{\mathrm{c}} = 3316$, where the sum now runs only over the cells.

### 3 Running the simulations

The simulations are run by numerically solving the equations of motion for the cells and collagen with a time-step size of $0.2\,\mathrm{s}$ using a modified version of the software package LAMMPS[3]. We started with a relaxation of the initial cell distribution for $1\,\mathrm{h}$ and afterwards performed the main simulations for a period of $12\,\mathrm{h}$, which agrees with the duration for which the cells are observed in the experiments.

**Initial relaxation.** As the nucleus of a real cell is usually not exactly in the centre of the cell, the initial distribution of the spherical particles representing cells in our simulations involves some overlap between particles. Therefore, we perform a relaxation of the initial particle configuration before the main simulations are run. For this purpose, we set up a homogeneous collagen grid with $\rho_{\text{loc}} = \bar{\rho}_0$ that encompasses the cells. Grid particles that overlap with cells are removed. With this setup, the time-evolution is calculated to allow the cell positions to relax, which results in a slight growth of the initial cancer spheroid. Since at the beginning of the experiments, the cells do not push against the collagen, we reduce the spring constants of the collagen grid during the relaxation phase to zero. The spring constants $k_{\text{nn}}(\vec{r}, \vec{r}')$ and $k_{\text{nnn}}(\vec{r}, \vec{r}')$ are therefore replaced by the functions

$$k_{\text{nn,relax}}(\vec{r}, \vec{r}', t) = k_{\text{nn}}(\vec{r}, \vec{r}')\Big(1 - \frac{t}{1\,\text{h}}\Big), \tag{7}$$

$$k_{\text{nnn,relax}}(\vec{r}, \vec{r}', t) = k_{\text{nnn}}(\vec{r}, \vec{r}')\Big(1 - \frac{t}{1\,\text{h}}\Big). \tag{8}$$

This results in a cell configuration that is rather close to the initial distribution obtained from the experiments and where the cells' positions are fully relaxed. During the relaxation, the radius of the cells is kept constant at $R_0$.

Next, the relaxed cells are placed in a collagen grid that corresponds to the initial collagen distribution in the experiments. Grid particles that overlap with the cells are removed. This ensures that the cells exert no stress on the collagen at the start of the main simulations. The obtained configuration of cancer cells and collagen is used as initial condition for the main simulations.

**Main simulations.** In the experiments, the outer layer of cells in contact with collagen and, with a slight time delay, a few inner layers within the cancer aggregate have been seen to inflate, starting at $t_0 = 6\,\text{h}$ (see Supplementary Figure 3). To reproduce this behaviour, we change the cells' radii as functions of their positions and time. For the first $6\,\mathrm{h}$ after the start of the main simulations, all cells have their initial radius $R_0$. Then, cells with a distance smaller than $d_{\mathrm{crit}}(t)$ to the nearest collagen particle start to inflate. The initial critical distance is $d_{\mathrm{crit}}(0) = 3R_0$ and this distance linearly grows with time so that it increases by $R_0$ within $1\,\mathrm{h}$:

$$d_{\mathrm{crit}}(t) = R_0\Big(3 + \frac{t}{1\,\mathrm{h}}\Big). \tag{9}$$

When $d_{\mathrm{crit}}$ has increased by $0.2R_0$, which occurs within $12\,\mathrm{min}$, the cell-collagen distance is determined again for each cell and for those cells, where the distance is below $d_{\mathrm{crit}}(t)$, inflation is started. An inflating cell linearly increases its radius by $R_0$ over the course of $2\,\mathrm{h}$ so that the radius of a cell that starts inflation at time $t_0$ is given by the function

$$R(t) = \begin{cases} R_0, & \text{if } t \leq t_0, \\ R_0 + R_0\frac{t-t_0}{2\,\mathrm{h}}, & \text{if } t_0 < t \leq t_0 + 2\,\mathrm{h}, \\ 2R_0, & \text{else.} \end{cases} \tag{10}$$

**Supplementary Videos**

Supplementary Video 1: **HeLa aggregates collectively burst in 0.5 mg ml$^{-1}$ collagen matrix.**

Bursts in LCC (0.5 mg ml$^{-1}$) and non-invasive behaviour in HCC (2.5 mg ml$^{-1}$).

Supplementary Video 2: **Nuclei tracking.**

Tracks of nuclei over 12 h in LCC and HCC.

Supplementary Video 3: **PAA beads as local pressure sensors.**

Aggregates containing elastic PAA beads as pressure sensors in LCC. Deformation of beads until bursts (zoom).

Supplementary Video 4: **Pushing forces exerted by aggregates on PAA gels.**

Sandwich experiment with deformation of the bottom PAA gel as aggregate pressure rises in confinement. Blue line as reference for the interface between aggregate and bottom gel.

Supplementary Video 5: **Collagen pockets - least mechanical resistance.**

Bursts of cells into collagen pockets in LCC.
